# Rapid GeneXpert surveillance of influenza A virus in seabirds and the environment provides early warning for wildlife health in Aotearoa New Zealand

**DOI:** 10.64898/2026.03.23.713605

**Authors:** Lia Heremia, Hoani Langsbury, Jackson Treece, Allison K Miller, Stephanie J Waller, James Ussher, Libby Manning, Christine Cleave, Zach Barford, Laura Findlay, Kate Cameron, David Micheal, Aliguna Aliguna, Tabitha Mason, Bella O’Connor, Steven G Badman, Neil J Gemmell, Jemma L Geoghegan, Jo-Ann L Stanton

**Affiliations:** Department of Microbiology and Immunology, University of Otago, Dunedin 9016, Aotearoa New Zealand; Royal Albatross Centre, Harington Point 9077, Aotearoa New Zealand; Department of Anatomy, University of Otago, Dunedin, Aotearoa New Zealand; Cepheid Holdings Pty Ltd, North Ryde, NSW 2113, Australia; Te Tātai Hauora o Hine National Centre for Women’s Health Research Aotearoa, Victoria University Wellington, Wellington 6140, Aotearoa New Zealand

**Keywords:** Avian influenza A virus, Highly pathogenic avian influenza (HPAI), Point-of-care diagnostics, GeneXpert, Environmental surveillance, Wildlife sentinel systems

## Abstract

The global expansion of highly pathogenic avian influenza (HPAI) virus A(H5N1) underscores the need for rapid surveillance at high-risk wildlife interfaces. Taiaroa Head (45.7828° S, 170.7333° E) in the South Island of Aotearoa New Zealand hosts a plethora of aquatic wildlife including a large red-billed gull (*Chroicocephalus novaehollandiae scopulinus*) colony as well as the only mainland breeding colony of northern royal albatross (*Diomedea sanfordi*). The Royal Albatross Centre is also a major nature tourism destination, attracting tens of thousands of visitors annually, thereby creating a dense ecological and human-wildlife interface vulnerable to viral incursion. We evaluated the GeneXpert II platform using the Xpert® Xpress Flu/RSV cartridge as a field-deployable tool for avian influenza virus detection in environmental and wildlife-associated samples. The assay detected synthetic influenza A viral RNA and multiple endemic low pathogenic avian influenza virus subtypes (A(H3N8), A(H1N9), A(H5N2) and A(H7N7)) circulating in New Zealand birds. Influenza A virus was reliably identified in spiked environmental water samples with no consistent PCR inhibition as well as naturally occurring avian influenza virus in duck pond water. Field deployment demonstrated that the system could be operated by non-laboratory personnel with minimal training in a non-clinical setting. This study establishes the feasibility of near-real-time environmental monitoring. Repurposing clinical cartridge-based point-of-care diagnostics offers a practical early warning approach for avian influenza virus surveillance at ecologically and economically significant locations.

## Introduction

Much of Aotearoa New Zealand’s iconic fauna is avian, reflecting an evolutionary history shaped by the near absence of terrestrial mammals (1). Birds have diversified to occupy ecological roles filled by mammals elsewhere, giving New Zealand a unique place in global natural history (2). Aquatic and coastal birds comprise a substantial component of this fauna, with many species undertaking long-distance migrations that bring them into close contact with avifauna from multiple regions (3). These migratory connections create pathways for the introduction and dissemination of infectious diseases (4). The ongoing global spread of highly pathogenic avian influenza (HPAI) virus subtype A(H5N1) clade 2.3.4.4b, combined with natural wildlife movements, places New Zealand at tangible risk of viral incursion (5,6).

The red-billed gull (Tarāpunga; *Chroicocephalus novaehollandiae scopulinus*) colony at Taiaroa Head, at the mouth of Otago Harbour, Dunedin (45.7828° S, 170.7333° E), is one of the largest in New Zealand, with more than 30,000 nesting birds (7). The colony sits adjacent to the internationally renowned northern royal albatross (Toroa; *Diomedea sanfordi*) breeding grounds, the only mainland albatross colony and a focal point of a major tourism operation (8). The headland also supports multiple other wildlife species, including little blue penguins (Kororā; *Eudyptula minor*), cormorants (Matapo; *Leucocarbo chalconotus*, Kawau Tikitiki; *Phalacrocorax punctatus* and Kawaupaka; *Microcarbo melanoleucos*), royal spoonbills (Kōtuku Ngutupapa; *Platalea regia*) and New Zealand fur seals (Kekeno; *Arctocephalus forsteri*) (9). Some of these species prey on other birds making them potentially susceptible to viral transmission (10-12). These overlapping populations create a complex ecological interface linking migratory birds, resident wildlife, marine mammals and humans through wildlife management and nature tourism. This convergence raises concerns for both conservation and public health, while also posing potential risks to an economically significant tourism asset in New Zealand’s South Island (13).

Rapid detection of HPAI virus at such high-risk interfaces is critical for timely management responses (14). The GeneXpert System (Cepheid) is a cartridge-based, point-of-care molecular diagnostic platform widely used for real-time detection of infectious diseases in human clinical settings (15-18) and has been shown to detect cultured A(H5N1) virus derived from cattle (19). Nevertheless, its performance in detecting avian influenza viruses from complex environmental and wildlife-derived samples, and its suitability as a field-deployable platform in remote settings with non-technical operators, has not yet been established. Each sealed cartridge integrates sample processing, PCR amplification and detection, enabling safe use by non-specialist users (20). Of particular relevance to this study, GeneXpert influenza A virus assays are not subtype-specific and should, in principle, detect a broad range of influenza A viruses, including those hosted by birds in New Zealand.

Early warning of an HPAI virus incursion would enable rapid implementation of containment and mitigation strategies (14). Given its size, density and ecological connectivity, sea birds present at Taiaroa Head represent an ideal sentinel system. However, effective on-site surveillance has been constrained by the lack of a rapid, accurate diagnostic method deployable in the field. This study evaluates whether the GeneXpert platform can fill this gap and support real-time avian influenza virus surveillance in a wildlife sentinel context.

## Methods

### GeneXpert II platform and assay conditions

All experiments were performed using a GeneXpert II instrument (Cepheid, USA) running GeneXpert® DX System Software version 6.5 and assay definition Xpert® Xpress Flu-RSV version 6, in conjunction with Xpert® Xpress Flu/RSV cartridges (XPRSFLU/RSV-CE-10; Cepheid, USA), according to the manufacturer’s instructions (20). The Sample Processing Control (SPC) is a non-target nucleic acid included in the cartridge that co-extracts and amplifies with sample nucleic acids, verifying effective sample processing, nucleic acid integrity, appropriate PCR conditions and the absence of significant inhibitors. Failure to meet SPC criteria in analyte-negative samples results in an invalid test outcome. All cartridges used in this study were from a single manufacturing lot (Kit Lot 10011442408; Reagent Lot ID: 16903) to ensure consistency of the internal SPC across experiments. The SPC was used to standardise assay performance and enable comparison across sample types and experimental conditions.

The GeneXpert platform performs automated nucleic acid extraction followed by a multiplex real-time reverse transcriptase polymerase chain reaction (RT-PCR) within a single cartridge, allowing simultaneous detection of *Influenza A virus, Influenza B virus* and *Respiratory syncytial virus* (RSV). The Xpert® Xpress Flu/RSV assay includes two independent target pools for *Influenza A virus* reported in the GeneXpert system as ‘Flu A 1’ and ‘Flu A 2’. Both pools use primers and probes that detect unique sequences in the genes encoding the matrix proteins (M1 and M2), polymerase basic 2 (PB2) and polymerase acidic (PA) (21). At least one target pool must be detected for a sample to be reported as positive for *Influenza A virus* (22). Results were classified as positive, negative or indeterminate (Invalid, Error, No Result), and the associated cycle threshold (Ct) and reaction endpoint (EndPt) fluorescence values were recorded in the GeneXpert system database.

For each run, 300 µl of sample, comprising diluent controls, environmental DNA (eDNA) samples, guano suspensions or raw fluid samples, was added directly to the cartridge using the transfer pipette supplied with the kit. Cartridges were loaded into the GeneXpert II instrument and processed immediately following sample addition.

### Spiked sample experiments

A range of diluents and environmental samples were evaluated, including samples spiked with known influenza A virus subtypes. These control samples included synthetic influenza A virus RNA, RNA derived from environmental and seabird metatranscriptomes with detected avian influenza virus (6,23), as well as RNA from viral isolates previously sampled from mallard ducks (5) (**Table 1**).

**Table 1.** Isolated RNA containing known avian influenza viruses for spiked experiments with GeneXpert.

| Subtype or gene | GenBank accession | Source/host | Method of identification | Experiment | Citation (if applicable) |
| --- | --- | --- | --- | --- | --- |
| M2 | PZ167570 | Synthetic | – | Spiked in PBS | New Zealand RNA Platform Research Platform |
| A(H3N8) | PZ167563-PZ167569 | Mallard duck, faeces | RNA sequencing | Spiked in PBS | Lia Heremia, unpublished data |
| A(H5N2) | MH231588 | Mallard duck, cloacal/oral swab | Virus isolation and genome sequencing | Spiked in PBS | New Zealand Ministry for Primary Industries |
| A(H7N7) | CY061618.1-CY061625.1 | Mallard duck, cloacal/oral swab | Virus isolation and genome sequencing | Spiked in PBS | New Zealand Ministry for Primary Industries |
| A(H1N9) | PQ358076-PQ358083 | Red knots, cloacal/oral swab | RNA sequencing | Spiked in PBS | Waller et al. 2025 |
| A(H3N2) | NC_007366-NC_007373 | Synthetic | – | Spiked in environmental water samples | Twist Bioscience |

Three standard diluents (PCR-grade water, 1x TE and 1x phosphate-buffered saline (PBS) (Invitrogen™)) and two routinely used RNA stabilisation buffers (RNALater (ThermoFisher AM7020) and RNA/DNA Shield (Zymo Research, R1100)) were tested for background impact on the Xpert® Xpress Flu/RSV cartridge chemistry. Four replicates of 300 µl for each diluent were tested and the SPC compared.

To test whether GeneXpert could detect a range of avian influenza virus subtypes, approximately 1,000 copies of influenza A virus RNA were spiked into 300 µl of 1× PBS for samples with known RNA concentrations (**Table 1**). This input exceeded the reported limit of detection of the Xpert® Xpress Flu/RSV assay (300 copies), ensuring detectable signal if amplification occurred. For samples where RNA concentration was unavailable or where RNA was derived from mixed environmental extracts, 2 µl of a 1:10 dilution of the stock RNA was added to 300 µl PBS. All samples were tested in duplicate to conserve material.

### Sample types and collection

Two sample types were used by staff at the Royal Albatross Centre for testing. First, dry guano samples were collected by scraping surface material with a sterile wooden applicator and resuspending the material in 5 ml of 1× PBS with vigorous shaking to homogenise the sample. Second, water samples were collected from standing water sources and birdbaths located at Taiaroa Head. Spiking experiments were conducted on ten standing water samples collected from the grounds of the Royal Albatross Centre which were spiked with 1,000 copies of synthetic A(H3N2) RNA (Twist Bioscience, **Table 1**) into 300 µl of undiluted sample.

To expand the environmental sample types tested, we also collected a range of samples from the Dunedin Botanic Gardens, Dunedin, which hosts a high density of mallard ducks and where we have previously detected avian influenza viruses via PCR and RNA sequencing (23). Water, sediment, and fresh duck faeces were collected. Water was filtered actively and passively using two methods. Active filters consisted of Wilderlab (https://wilderlab.co/) Wetland kits and passive filters consisted of sterile sponges that were deployed on floats overnight for ∼16 hours. Approximately 30ml of submerged sediment was collected from ∼10 cm below the surface. Fresh faecal samples were obtained using a sterile flock swab and placed into RNA/DNA Shield (Zymo Research). Using the transfer pipette supplied, 300 µl of either the resuspended sample or undiluted water sample was added directly to the GeneXpert cartridge for testing.

### Field deployment and user operation

The GeneXpert II instrument was deployed in a room adjacent to the public wildlife observation facility operated by the Royal Albatross Centre at Taiaroa Head (**Figure 1**). Centre staff received training in sample collection and operation of the GeneXpert system prior to deployment.

**Figure 1.**
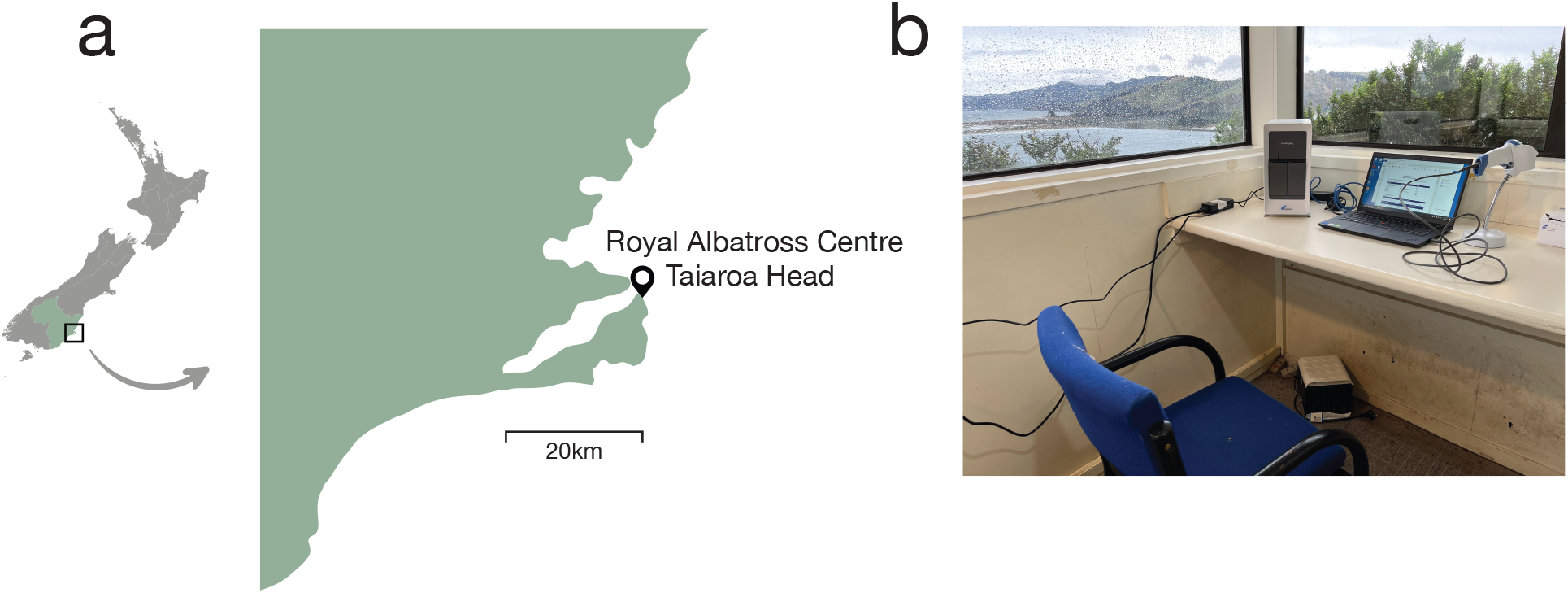
(a) Map of New Zealand with an inset of the Otago region (green) showing the Otago Peninsula in Dunedin, highlighting the location of the Royal Albatross Centre at Taiaroa Head (45.7828° S, 170.7333° E). (b) GeneXpert set up at the Royal Albatross Centre.

Participating staff had no prior formal laboratory training. The instrument was installed on 9 May 2025 and remained in continuous operation until 24 November 2025. Following removal of the instrument, participating staff were invited to provide feedback on their experiences using the GeneXpert through informal interviews.

## Results

### Assay performance across sample diluents

The effect of sample diluent on assay performance was assessed by examining the Ct and EndPt fluorescence values of the manufacturer-supplied SPC. The SPC is a positive PCR target added during cartridge manufacture and serves as an internal indicator of assay integrity. SPC failure indicates inhibition of on-board chemistry (20). All cartridges used in this analysis were from a single manufacturing lot, ensuring comparability across runs.

Five diluents were evaluated, with SPC Ct and EndPt values calculated from four replicate runs per diluent. Ct values differed among preservation treatments (one-way ANOVA, F(4,15) = 4.69, p = 0.012) (**Figure 2, Table S1**). Tukey’s honestly significant difference (HSD) test showed that RNAlater samples had significantly higher Ct values than PBS (p = 0.014), water (p = 0.018), and RNA/DNA Shield (p = 0.049), with no other significant pairwise differences. EndPt values also differed among diluents (F(4,15) = 7.86, p = 0.0013), with RNAlater producing significantly lower EndPt values than PBS (p = 0.001), water (p = 0.003) and TE (p = 0.009), indicating reduced amplification performance in RNAlater-preserved samples relative to the other buffers.

**Figure 2.**
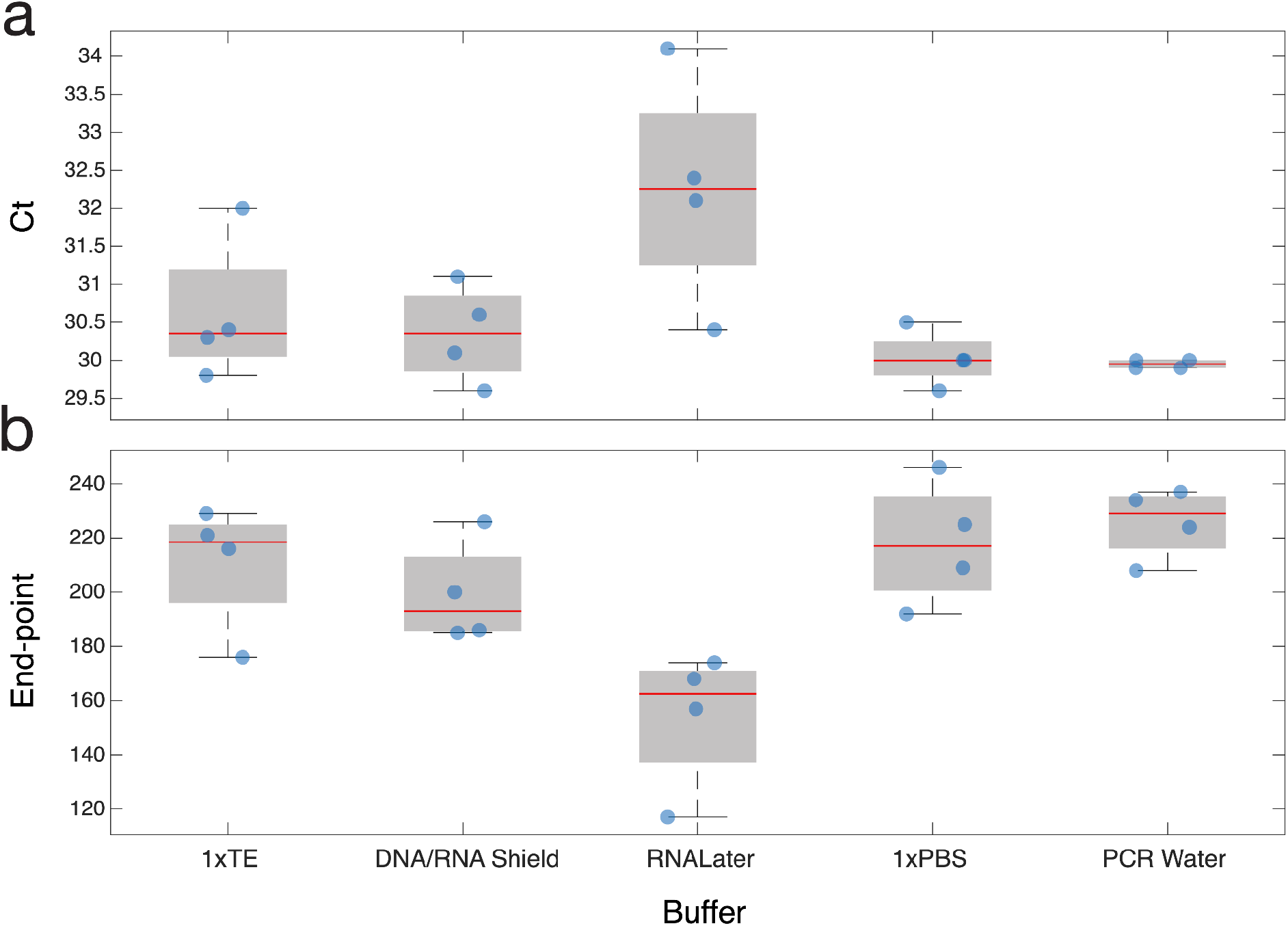
(a) SPC Cycle threshold (Ct) and (b) End-point values for five diluents tested with the GeneXpert. The central line (red) indicates the median, the grey box represents the interquartile range (IQR; 25th–75th percentiles), whiskers extend to 1.5× the IQR and points beyond this range shown as outliers. Raw data are superimposed for clarity.

### Detection of endemic avian influenza A viral subtypes

Compatibility of the Xpert® Xpress Flu/RSV assay with endemic avian influenza A virus strains circulating in New Zealand was evaluated using extracted viral RNA and synthetic controls. RNA from avian influenza virus isolated from environmental and avian samples, as well as synthetic influenza A virus, were tested in duplicate (**Table S2**). Tested material included A(H3N8) derived from mallard duck (*Anas platyrhynchos*) faeces, A(H1N9) from cloacal swabs collected from bar-tailed godwits (*Limosa lapponica*) (6), A(H5N2) and A(H7N7) derived from viral isolates detected in mallard ducks (*Anas platyrhynchos*), along with a synthetic influenza A virus matrix protein 2 (M2) gene (RNA platform control, G-Block AI_002). The assay successfully detected the synthetic M2 RNA as well as A(H3N8), A(H5N2), A(H7N7) and A(H1N9) obtained from a range of aquatic birds (**Figure 3**). Both Flu A 1 and Flu A 2 targets detected all influenza A virus subtypes. Ct values differed relative to the SPC (mean Ct: Flu A 1 = 31.3, Flu A 2 = 32.6, SPC = 30.4), with Flu A 2 significantly higher than the SPC (paired t-test, p = 0.008), while Flu A 1 did not differ significantly (p = 0.28). In contrast, EndPt fluorescence was significantly higher for both Flu A targets compared with the SPC (mean EndPt: Flu A 1 = 574.7, Flu A 2 = 299.1, SPC = 185.0; paired t-tests p ≈ 0.002 for both), indicating strong target-specific amplification and demonstrating that the GeneXpert Xpress Flu/RSV cartridge can detect influenza A virus strains circulating in wild New Zealand bird populations (**Figure 3**).

**Figure 3.**
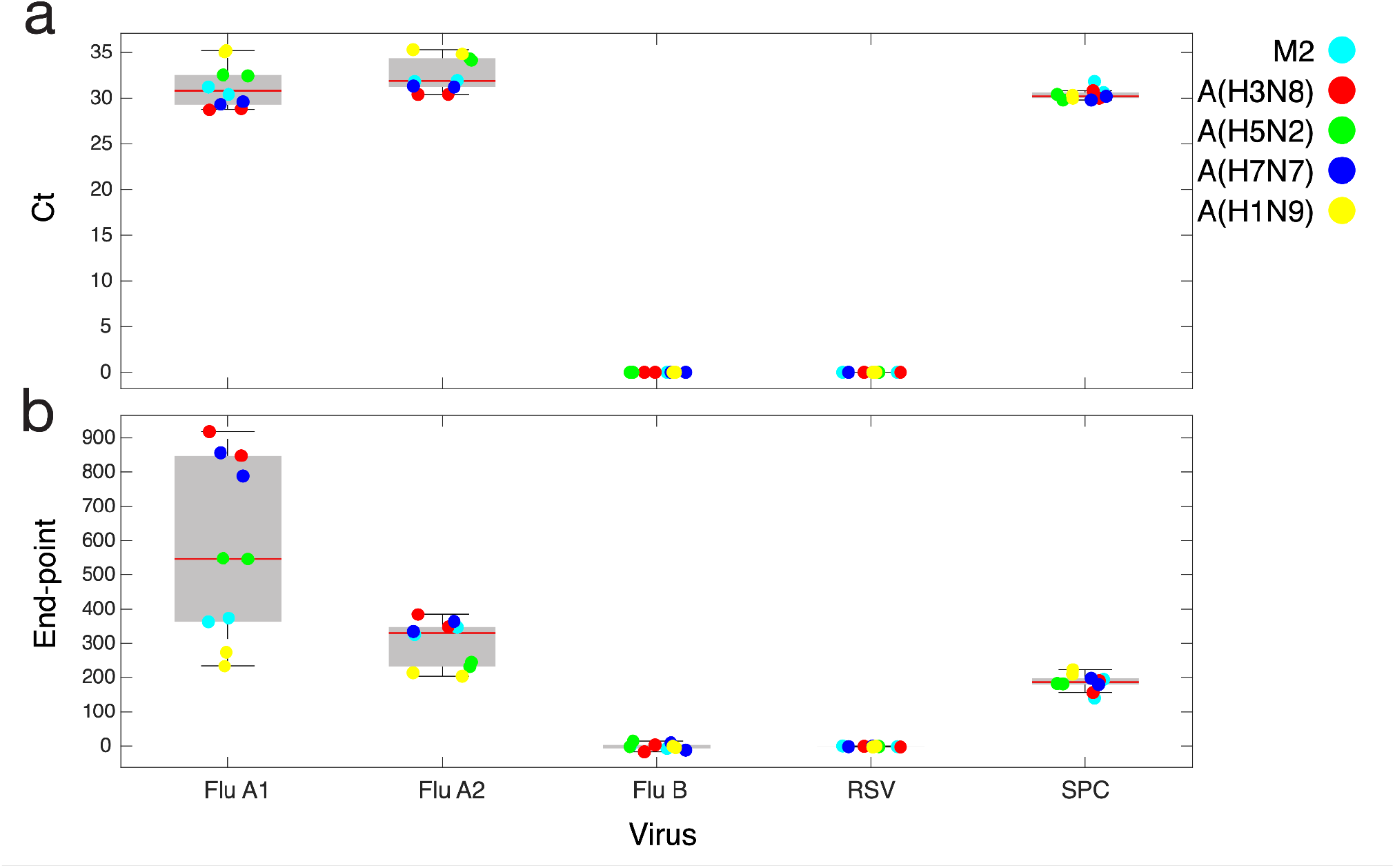
(a) Cycle threshold (Ct) and (b) End-point values for spiked experiments with A(H3N8) (red) derived from mallard duck (*Anas platyrhynchos*) faeces, A(H1N9) (yellow) from cloacal swabs collected from bar-tailed godwits (*Limosa lapponica*) (6), A(H5N2) (green) and A(H7N7) (blue) derived from viral isolates detected in mallard ducks (*Anas platyrhynchos*), along with a synthetic matrix protein 2 (M2) gene (cyan). The central line (red) indicates the median, the grey box represents the interquartile range (IQR; 25th–75th percentiles), whiskers extend to 1.5× the IQR and points beyond this range shown as outliers. Raw data are superimposed for clarity.

### Detection of influenza A virus in spiked environmental water samples

To assess assay performance in complex environmental samples, ten standing water samples collected from the grounds of the Royal Albatross Centre (**Figure S1**) were spiked with synthetic A(H3N2) RNA (Twist Bioscience) into 300 µl of undiluted sample. Synthetic A(H3N2) was detected across all samples by both Flu A targets (**Figure 4, Table S3**). Ct values were significantly higher for Flu A 1 (34.0 ± 1.5) and Flu A 2 (35.8 ± 1.1) than for the SPC (30.6 ± 0.6) (paired t-tests: Flu A 1 vs SPC, p < 0.001; Flu A 2 vs SPC, p < 0.001). EndPt fluorescence values also differed from the control, with Flu A 1 producing significantly higher signals than the SPC (p < 0.001), while Flu A 2 EndPt did not differ significantly from the SPC (p = 0.11).

**Figure 4.**
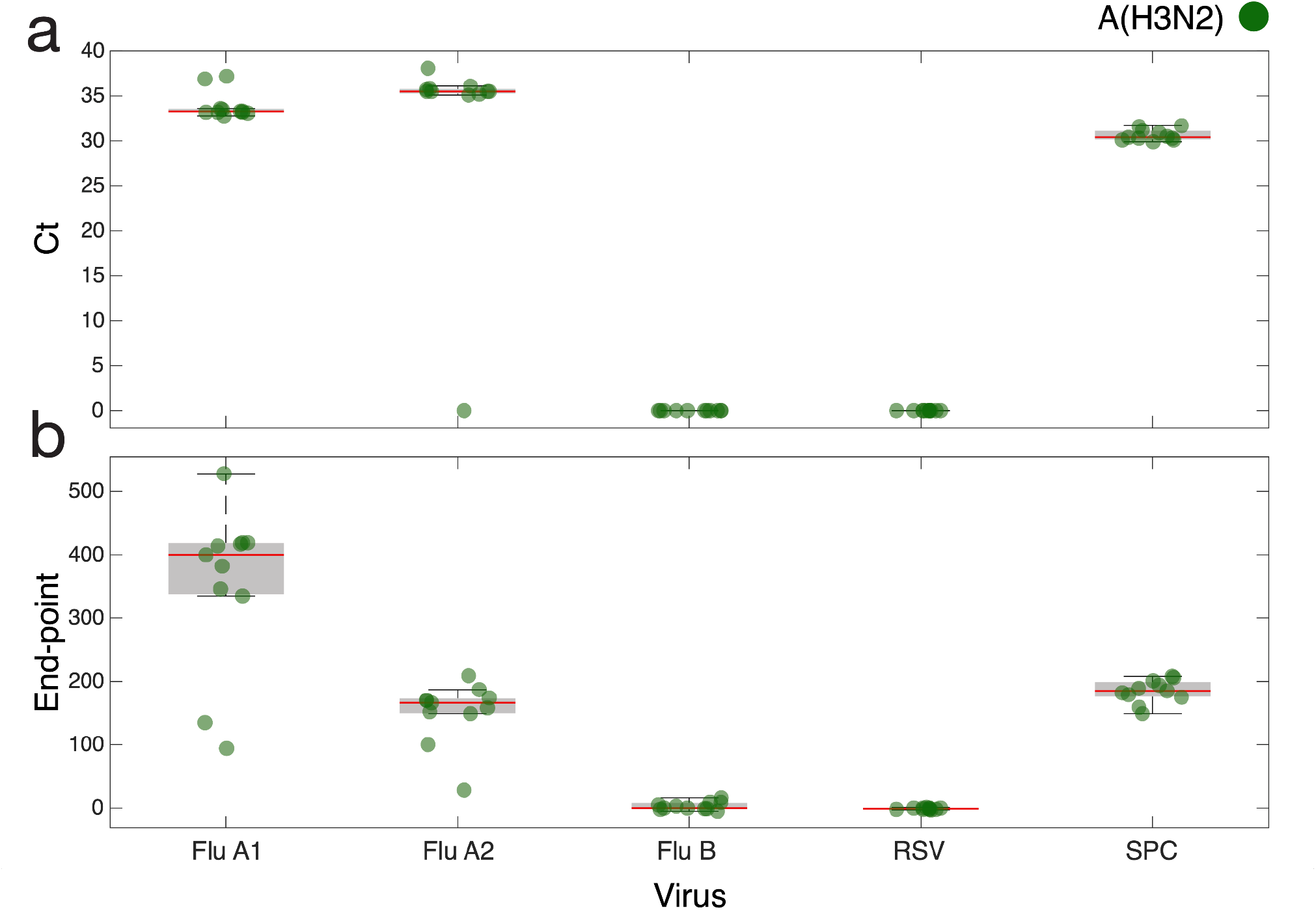
(a) Cycle threshold (Ct) and (b) End-point values for ten standing water samples collected from the grounds of the Royal Albatross Centre spiked with 1,000 copies of synthetic A(H3N2) RNA. The central line (red) indicates the median, the grey box represents the interquartile range (IQR; 25th–75th percentiles), whiskers extend to 1.5× the IQR and points beyond this range shown as outliers. Raw data are superimposed for clarity.

One sample (Car Park #5) was visibly discoloured, suggesting a high contaminant load (**Figure S1**). This sample exhibited higher Ct values than other spiked water samples (mean Ct 37.05 for Flu A 1), with Flu A 2 detected in only one of two replicates (Ct 38.1) (**Table S3**). Notably, SPC Ct values for this sample (mean 30.25) were comparable to those of other samples, indicating intact assay chemistry. These results suggest partial degradation of spiked influenza viral RNA, potentially due to environmental RNase activity, rather than PCR inhibition.

### Field detection of influenza A virus in wildlife-associated water sources

A range of environmental samples were also collected from a large urban duck pond in the Dunedin Botanic Garden, where waterfowl are known reservoirs of endemic avian influenza viruses (23). Samples, including fresh faeces, passive filtered water, standing water and sediment, from three sites around the pond were tested. Influenza A virus positive samples were detected at two locations in standing pond water samples, but not in passive water, sediment or faecal samples (**Figure 5, Table S4**). Specifically, influenza A virus was detected in 10/15 samples by Flu A 1 and 4/15 by Flu A 2, with Ct values for positive samples ranging from 34.2-38.7 and 36.7-39.3, respectively, while the SPC remained consistent (31.5 ± 1.9). Ct values for positive samples were significantly higher than the SPC (Flu A 1 paired t-test p < 0.001; Flu A 2 p < 0.001). EndPt fluorescence values for positive samples were higher for Flu A 1 than the SPC (p = 0.03), whereas Flu A 2 EndPt values did not differ significantly from the SPC (p = 0.12). These findings demonstrate that the GeneXpert II platform coupled with the Xpert® Xpress Flu/RSV assay can detect naturally occurring influenza A virus from environmental water samples.

**Figure 5.**
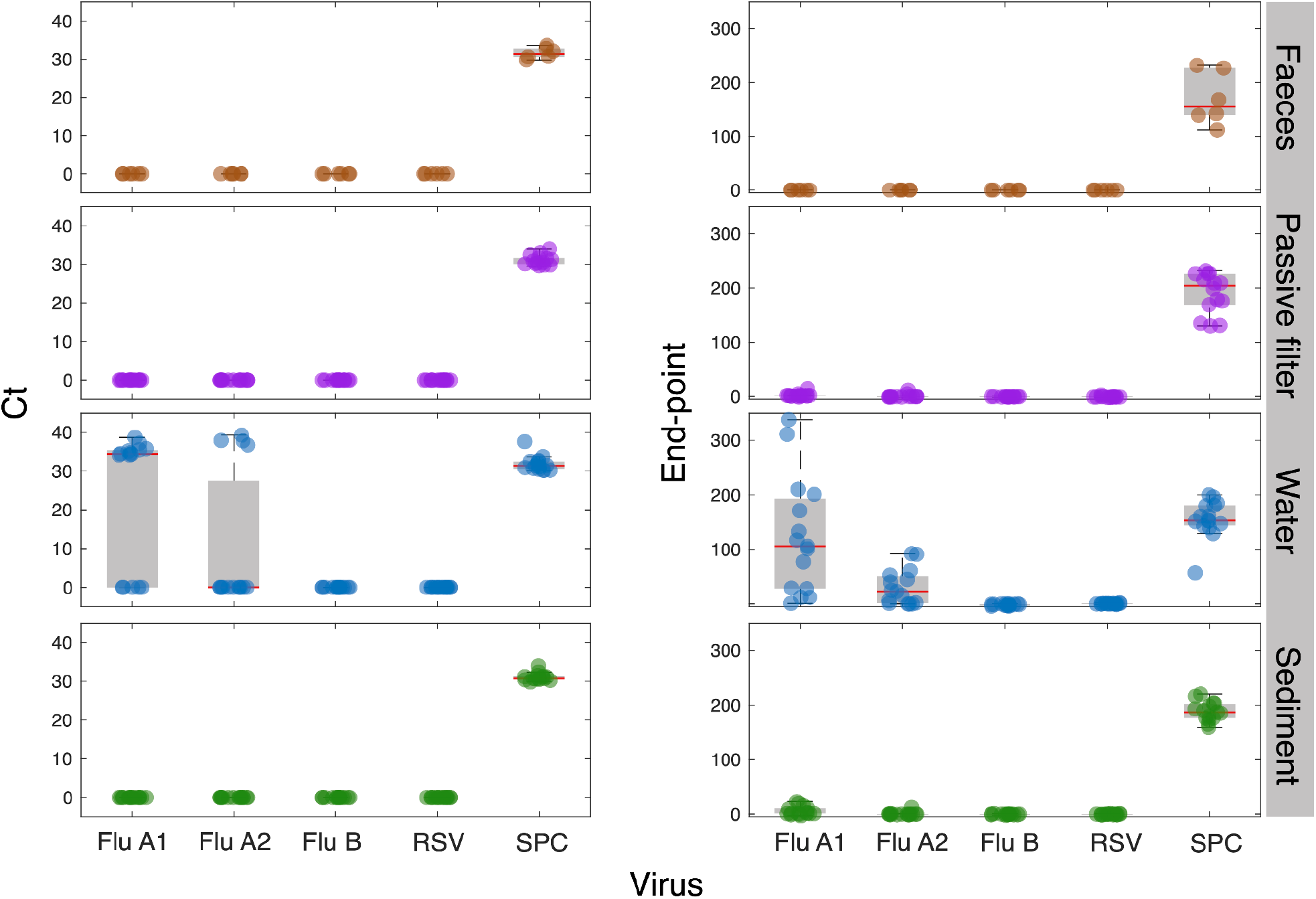
(left) Cycle threshold (Ct) and (right) End-point values for a range of different environmental samples including fresh faeces, passive filtered water, standing water and sediment obtained from the pond in the Dunedin Botanical Gardens. The central line (red) indicates the median, the grey box represents the interquartile range (IQR; 25th–75th percentiles), whiskers extend to 1.5× the IQR and points beyond this range shown as outliers. Raw data are superimposed for clarity.

### Field deployment and sentinel surveillance at Taiaroa Head

Based on laboratory and environmental validation results, an implementation study was undertaken at Taiaroa Head, Otago Peninsula. The GeneXpert II instrument was installed in an unheated, uninsulated room with mains power and plumbed water at the Royal Albatross Centre, overlooking Otago Harbour (**Figure 1**). Over the course of the study, a portable heater was placed in the room as ambient temperatures fell below the operational limits of the instrument. Staff received a single training session and had access to remote technical support as needed. During the deployment period, surveillance focused on non-invasive sampling, including guano scrapings and water samples collected from temporary birdbaths installed at the site. No birds were captured or handled. Influenza A virus was not detected in any samples during the deployment period (**Table S5**).

The GeneXpert II experienced increased run errors following placement at the Taiaroa Head site. Most errors (21/22) were associated with syringe pressure (Error 2008: Syringe pressure reading of 100.1 PSI exceeds the protocol limit of 100.0 PSI). Errors were associated with processing both guano and water samples (9 guano and 12 water samples) and hypothesised to result from particulates in the sample. This suggests adding a filtration step or allowing samples to settle during sample preparation may be helpful. However, the instrument underwent double module replacement following removal from the Taiaroa Head facility. This was undertaken on advice from Cepheid engineers. An error log for the full project is given in **Table S6**.

Following removal of the GeneXpert II instrument from the Royal Albatross Centre, staff participated in small-group interviews to share their experiences using the system. A qualitative review of these discussions identified several recurring themes, including ease of use, perceived value for wildlife health surveillance, and the potential role of point-of-care molecular diagnostics in conservation settings (**Table 2**).

**Table 2.** Summary of staff perspectives on the implementation of the GeneXpert II system at the Royal Albatross Centre.

| Theme | Key observations | Implications for future surveillance |
| --- | --- | --- |
| **Ease of use and workflow integration** | Staff integrated testing into daily duties by scheduling runs alongside guided tours, sharing responsibilities across multiple operators, and allocating specific time blocks for testing. Technical confidence improved rapidly and the system was eventually viewed as “not a stressor”. | Training multiple operators and embedding testing into existing workflows can facilitate adoption of point-of-care diagnostics in conservation settings. |
| **Operational guidance and SOPs** | Manuals and site-specific SOPs were used for procedural reference. Staff recommended expanded SOPs covering environmental sampling (e.g. guano and birdbath collection) and guidance on sampling frequency and distribution. | Clear SOPs for both instrument operation and sample collection are essential for consistent wildlife disease surveillance. |
| **Environmental sampling considerations** | Birdbaths were viewed as an effective way to sample the wider bird population, while guano sampling was straightforward during peak nesting. Sampling of rainwater tanks was also feasible. | Environmental sampling strategies require further optimisation to determine ideal sampling locations, timing, and frequency. |
| **Infrastructure and instrument placement** | The GeneXpert instrument was located in a remote, non-temperature-controlled room, increasing time required for runs and creating logistical barriers. | Instrument placement in accessible, temperature-controlled environments would improve operational efficiency. |
| **Technical challenges** | Pressure errors occasionally caused run termination, leading some staff to run duplicate samples. Allowing samples to settle after mixing appeared to reduce errors. | Environmental sample preparation protocols may need refinement to reduce cartridge pressure errors. |
| **Training and laboratory practices** | Staff were primarily trained in ecological fieldwork rather than laboratory practice. Some conventions such as frequent glove changes required clarification. | Explicit operational guidelines and training in contamination control and reagent management are needed. |
| **Value for surveillance and communication** | Staff viewed GeneXpert as useful for early detection of infection and enabling rapid response. The system also supported public engagement by demonstrating active disease monitoring. | Point-of-care diagnostics could enhance both wildlife disease surveillance and public communication at ecotourism sites. |

Testing was successfully incorporated into routine duties by allocating time for runs within work schedules, coordinating multiple operators and aligning testing with guided tours (**Table 2**). Technical confidence improved rapidly and the system was ultimately regarded as straightforward to operate. Manuals and site-specific Standard Operating Procedures (SOPs) were used when required, although staff recommended additional guidance for environmental sampling, including protocols for guano and birdbath sampling.

Environmental sampling approaches were considered practical. Birdbaths were viewed as an effective method for sampling the broader bird population, while guano collection was straightforward during periods of high nesting density. However, further work is required to determine optimal sampling locations and frequency (**Table 2**). Operational challenges were largely logistical, particularly the remote, non-temperature-controlled location of the instrument, which increased the time required to initiate and monitor runs. Despite these constraints, staff viewed the GeneXpert system positively and considered it a valuable tool for early pathogen detection and public engagement in wildlife health surveillance.

## Discussion

This study demonstrates that the GeneXpert II platform, using the Xpert® Xpress Flu/RSV assay, can be successfully applied to detect avian influenza viruses in environmentally relevant samples collected from a high-risk wildlife interface. At Taiaroa Head, red-billed gulls and other sea birds naturally aggregate around fresh water for bathing, creating predictable points of environmental fouling. In addition to temporary bathing pools established within the colony, faecal contamination of rainwater collection tanks at the visitor centre was observed, reflecting sustained interaction between wildlife, shared water sources and human infrastructure. These behaviours create plausible pathways for environmental accumulation of potential pathogens, including avian influenza virus, and support the rationale for water-based surveillance in this setting (24,25).

Samples collected from rainwater collection tanks, temporary bathing pools, resuspended guano and a range of pond-associated samples such as water, sediment and duck faeces were processed successfully using the GeneXpert II system, with no evidence of systematic PCR inhibition. Influenza A virus was readily detected in spiked samples, including those with visible organic contamination, and SPC performance remained stable across sample types. Positive influenza A virus detection was demonstrated in water samples collected from environments where low pathogenic avian influenza viruses were likely to occur naturally (26,23). These results indicate that GeneXpert II can tolerate complex environmental samples and generate interpretable results without extensive modification. However, an issue with particulates in environmental samples suggests it would be beneficial if it included an additional pre-processing step, such as filtration or a settling period prior to cartridge loading. Overall, the general system compatibility and ease of use of the GeneXpert II are key requirements for near-real-time surveillance in non-laboratory settings, where cold-chain access, specialist equipment and trained personnel may be limited.

This work was undertaken in response to the global threat of highly pathogenic A(H5N1) and the risk of its incursion into New Zealand. The Royal Albatross Centre sits at a nexus of wildlife and human interaction that presents a risk of HPAI introduction and spread. The viral subtypes tested in this work were chosen as they are already circulating in the New Zealand environment and presented an opportunity to rapidly determine the usefulness of GeneXpert System II in a real-world setting, as seen with non-spiked, naturally occurring environmental samples (**Figure 5**). The Xpress® Xpert Flu/RSV kit by design does not use the highly variable surface proteins, hemagglutinin (HA) or neuraminidase (NA), as specific markers for detection. Instead, it targets multiple conserved regions of the influenza A virus genome, improving robustness across diverse viral lineages. The assay incorporates two independent influenza A virus target channels (Flu A 1 and Flu A 2), each comprising multi-gene target pools designed to accommodate genomic variation among circulating strains. These features increase the likelihood of detecting genetically diverse influenza A viruses. Indeed, A(H5N1) has previously been shown to be successfully detected by this assay (19).

In conclusion, GeneXpert II was readily integrated into the operational workflows of the Royal Albatross Centre, despite initially being deployed in an unheated, non-laboratory environment and operated by staff without prior laboratory training. Minimal training was required, and the system performed as expected over several months of field deployment. The study established the feasibility of rapid, on-site molecular screening as part of an early warning tool. Future work should focus on optimising sample collection methods, expanding surveillance across diverse locations and evaluating assay performance against other environmental sample types. In addition, it would be valuable to assess whether integrating subtype- and clade-specific influenza A virus assays into a dedicated cartridge could enable rapid discrimination of high-consequence subtypes following initial detection. Consideration should also be given to how results are interpreted by non-expert users, as the lack of subtype specificity means positive detections could be mistakenly attributed to HPAI, highlighting the importance of clear interpretive guidance and adequate training in non-specialist settings. Overall, these findings support the use of point- of-care molecular diagnostics as a practical and viable addition to wildlife disease surveillance at critical conservation and human-wildlife interfaces.

## Data availability

Genetic data for influenza A virus RNA used in this study can be found under accession numbers listed in Table 1.

## Author contributions

Conceptualisation: JLS, HL, JLG, SGB, JU. Writing - originally draft: JLS, JLG, LH. Investigation: LH, JT, AKM, LM, CC, ZB, LF, KC, DM, AA, TM, BO’C, SW, JLS. Supervision: NJG, LM, LF, JLS.

Data curation: JLS. Formal analysis: JLG, LH, JLS. Reviewing and Editing: All.

## Acknowledgments

We thank the New Zealand Ministry for Primary Industries for providing avian influenza virus isolates for controlled spiked experiments.

