## Supplementary Figure 1 for "Rapid GeneXpert surveillance of influenza A virus in seabirds and the environment provides early warning for wildlife health in Aotearoa New Zealand"

**Supplementary Figure 1**. Water samples collected around Albatross Colony at Taiaroa Head between 25-27 Febrary 2025. All samples spiked with 1ul of H3N2 synthetic RNA @ ~1000cp/ul.

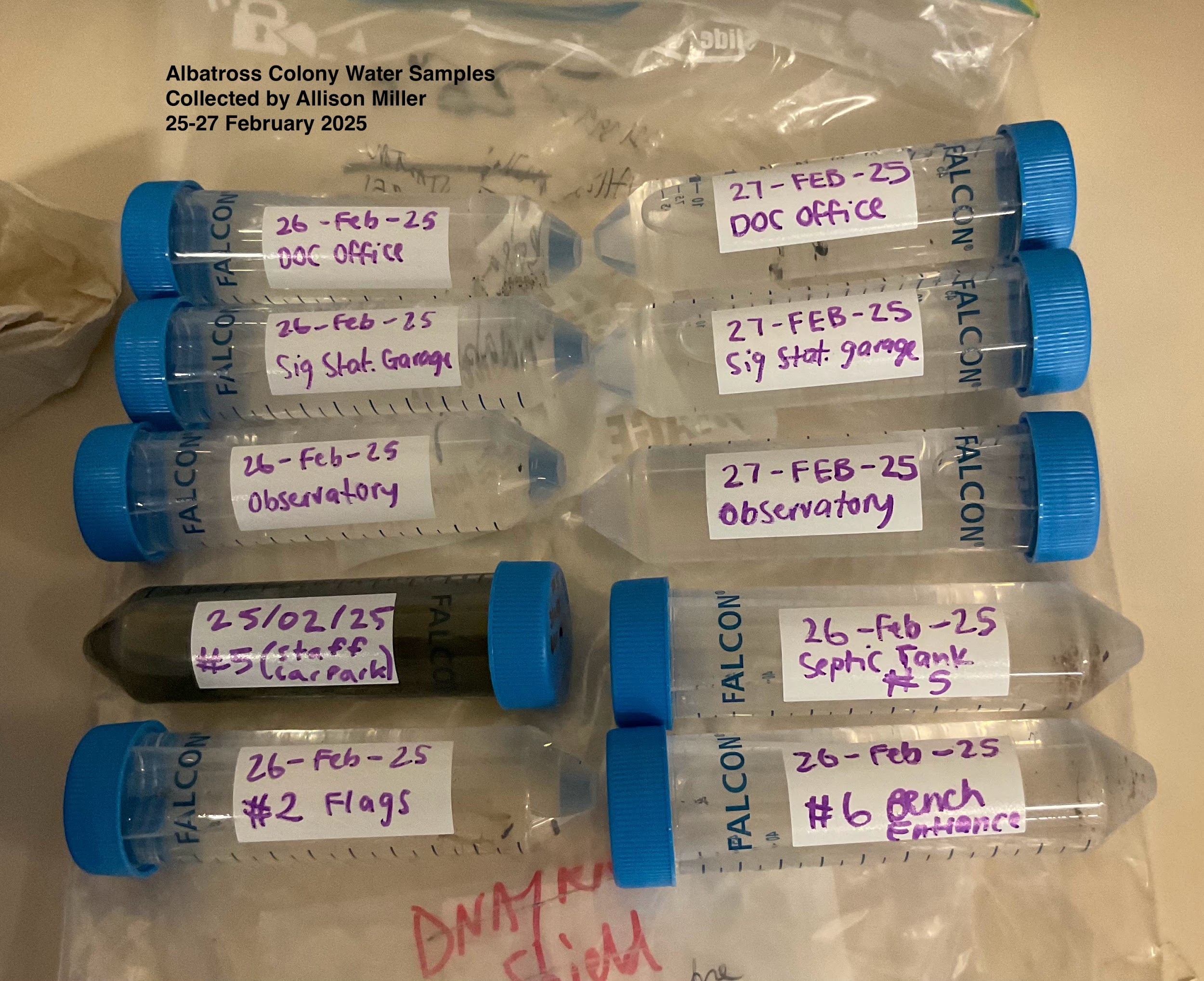
