## Supplementary Table 1 for "Rapid GeneXpert surveillance of influenza A virus in seabirds and the environment provides early warning for wildlife health in Aotearoa New Zealand"

**Supplementary Table 1**. GeneXpert optimisation with various sample collection buffers, showing the Ct and end-point of the Sample Processing Control (SPC) using Xpress Xpert Flu/RSV® cartridges, Lot Kit Lot 10011442408; Reagent Lot ID: 16903.

| Ct | 1x TE | DNA/RNA Shield | RNAlater | PCR Water | PBS |
| --- | --- | --- | --- | --- | --- |
|  | 29.8 | 29.6 | 32.4 | 29.6 | 29.9 |
|  | 30.4 | 31.1 | 32.1 | 30.5 | 30 |
|  | 32 | 30.1 | 30.4 | 30 | 30 |
|  | 30.3 | 30.6 | 34.1 | 30 | 29.9 |
| End-point | **1x TE** | **DNA/RNA Shield** | **RNAlater** | **PCR Water** | **PBS** |
|  | 229 | 226 | 168 | 246 | 237 |
|  | 216 | 185 | 157 | 192 | 208 |
|  | 176 | 200 | 174 | 225 | 224 |
|  | 221 | 186 | 117 | 209 | 234 |
