## Supplementary Table 2 for "Rapid GeneXpert surveillance of influenza A virus in seabirds and the environment provides early warning for wildlife health in Aotearoa New Zealand"

**Supplementary Table 2**. Duplicate spiking experiments with synthetic M2 gene as well as a range of known avian influenza viruses obtained from aquatic birds.

| Ct | M2_1 | M2_2 | H3N8_1 | H3N8_2 | H5N2_1 | H5N2_2 | H7N7_1 | H7N7_2 | H1N9_1 | H1N9_2 |
| --- | --- | --- | --- | --- | --- | --- | --- | --- | --- | --- |
| Flu A 1 | 31.2 | 30.4 | 28.8 | 28.7 | 32.4 | 32.5 | 29.3 | 29.6 | 35 | 35.2 |
| Flu A 2 | 31.9 | 31.8 | 30.4 | 30.4 | 34.1 | 34.3 | 31.2 | 31.3 | 34.8 | 35.3 |
| Flu B | 0 | 0 | 0 | 0 | 0 | 0 | 0 | 0 | 0 | 0 |
| RSV | 0 | 0 | 0 | 0 | 0 | 0 | 0 | 0 | 0 | 0 |
| SPC | 30.6 | 31.8 | 30.8 | 30 | 30.4 | 29.8 | 29.8 | 30.1 | 30.3 | 30 |
| End-point | M2_1 | M2_2 | H3N8_1 | H3N8_2 | H5N2_1 | H5N2_2 | H7N7_1 | H7N7_2 | H1N9_1 | H1N9_2 |
| Flu A 1 | 362 | 373 | 848 | 918 | 546 | 548 | 857 | 788 | 233 | 274 |
| Flu A 2 | 346 | 325 | 347 | 384 | 245 | 231 | 363 | 334 | 203 | 213 |
| Flu B | -7 | 0 | 3 | -17 | -2 | 14 | -12 | 9 | -5 | -1 |
| RSV | 0 | -2 | -3 | -1 | -1 | -2 | 0 | -2 | -1 | -3 |
| SPC | 194 | 140 | 156 | 190 | 182 | 181 | 197 | 179 | 209 | 222 |
