## Supplementary Table 3 for "Rapid GeneXpert surveillance of influenza A virus in seabirds and the environment provides early warning for wildlife health in Aotearoa New Zealand"

**Supplementary Table 3**. Water samples collected around Albatross Colony at Taiaroa Head between 25-27 Febrary 2025. All samples spiked with 1ul of H3N2 synthetic RNA @ ~1000cp/ul.

| **Ct** | **#5 staff car park 1** | **#5 staff car park 2** | **#2 flags** | **#6 bench entrance** | **#5 septic tank** | **observatory 1** | **observatory 2** | **sig stat garage 1** | **sig stat garage 2** | **DOC office 1** | **DOC office 2** |
| --- | --- | --- | --- | --- | --- | --- | --- | --- | --- | --- | --- |
| Flu A 1 | 36.9 | 37.2 | 33.3 | 33.2 | 33.1 | 33.6 | 33.2 | 33.2 | 33.5 | 32.8 | 33.3 |
| Flu A 2 | 38.1 | 0 | 35.5 | 35.5 | 35.5 | 36.1 | 35.5 | 35.2 | 35.7 | 35.1 | 35.8 |
| Flu B | 0 | 0 | 0 | 0 | 0 | 0 | 0 | 0 | 0 | 0 | 0 |
| RSV | 0 | 0 | 0 | 0 | 0 | 0 | 0 | 0 | 0 | 0 | 0 |
| SPC | 30.1 | 30.4 | 30.9 | 30.5 | 30.3 | 31.6 | 29.9 | 30.1 | 31.7 | 30.3 | 31.2 |
| **End-point** | **#5 staff car park 1** | **#5 staff car park 2** | **#2 flags** | **#6 bench entrance** | **#5 septic tank** | **observatory 1** | **observatory 2** | **sig stat garage 1** | **sig stat garage 2** | **DOC office 1** | **DOC office 2** |
| Flu A 1 | 135 | 94 | 417 | 400 | 419 | 346 | 414 | 419 | 382 | 528 | 335 |
| Flu A 2 | 100 | 28 | 166 | 174 | 158 | 149 | 170 | 187 | 170 | 209 | 152 |
| Flu B | 0 | 3 | -2 | 0 | 9 | -1 | 9 | -1 | 5 | -5 | 16 |
| RSV | 0 | 0 | -2 | 0 | -2 | 1 | -2 | 0 | -3 | -1 | -2 |
| SPC | 182 | 179 | 194 | 185 | 189 | 159 | 201 | 206 | 175 | 208 | 149 |
