## Supplementary Table 4 for "Rapid GeneXpert surveillance of influenza A virus in seabirds and the environment provides early warning for wildlife health in Aotearoa New Zealand"

**Supplementary Table 4**. GeneXpert runs performed by staff at the Royal Albatross Centre between July and October 2025.

| **Sample type** | **Date** | **Result** | **Module** | **SPC Ct** | **SPC EndPt** | **Flu A1 Ct** | **Flu A1 EndP** | **Flu A2 Ct** | **Flu A2 Endpt** |
| --- | --- | --- | --- | --- | --- | --- | --- | --- | --- |
| Guano | 02/09/2025 | Negative | B1 | 30.2 | 228 | 0 | -1 | 0 | -2 |
| Guano | 29/08/2025 | Negative | B1 | 29.9 | 253 | 0 | 0 | 0 | -3 |
| Guano | 31/08/2025 | Negative | B1 | 31.2 | 207 | 0 | 1 | 0 | -1 |
| Guano |  | Negative | B1 | 30 | 228 | 0 | 0 | 0 | -2 |
| Guano |  | Negative | B2 | 32.9 | 162 | 0 | -4 | 0 | -1 |
| Guano | 07/08/2025 | Negative | B2 | 31.3 | 188 | 0 | -1 | 0 | -1 |
| Guano | 08/09/2025 | Negative | B2 | 30.9 | 198 | 0 | 0 | 0 | -1 |
| Guano | 15/07/2025 | Negative | B1 | 31.1 | 164 | 0 | 0 | 0 | -1 |
| Guano | 16/09/2025 | Negative | B1 | 30.3 | 216 | 0 | 1 | 0 | -1 |
| Guano | 29/07/2025 | Negative | B1 | 33.1 | 173 | 0 | -2 | 0 | 0 |
| Guano | 26/07/2025 | Negative | B1 | 30.6 | 205 | 0 | -2 | 0 | -1 |
| Guano | 26/07/2025 | Negative | B2 | 31.6 | 200 | 0 | -2 | 0 | 0 |
| Guano |  | Negative | B2 | 34.9 | 116 | 0 | -2 | 0 | -1 |
| Guano | 02/10/2025 | Negative | B2 | 30.7 | 217 | 0 | 0 | 0 | -1 |
| Guano | 11/09/2025 | Negative | B2 | 33.3 | 160 | 0 | -1 | 0 | 0 |
| Guano | 09/09/2025 | Negative | B2 | 31.2 | 193 | 0 | -1 | 0 | 0 |
| Guano | 28/09/2025 | Negative | B2 | 32.5 | 169 | 0 | -1 | 0 | 0 |
| Guano | 14/08/2025 | Negative | B1 | 35.4 | 109 | 0 | 0 | 0 | -2 |
| Guano | 15/10/2025 | Negative | B2 | 32.5 | 163 | 0 | 22 | 0 | 37 |
| Guano | 28/08/2025 | Negative | B1 | 31.6 | 208 | 0 | 1 | 0 | -2 |
| Guano | 02/08/2025 | Negative | B2 | 31.1 | 204 | 0 | -1 | 0 | 0 |
| Guano | 18/08/2025 | Negative | B2 | 30.7 | 207 | 0 | -1 | 0 | 0 |
| Guano | 18/08/2025 | Negative | B1 | 30.1 | 219 | 0 | 1 | 0 | -2 |
| Guano |  | Negative | B1 | 30.2 | 243 | 0 | 1 | 0 | -2 |
| Guano |  | Negative | B2 | 31.5 | 186 | 0 | -1 | 0 | 0 |
| Bird bath / rain water | 28/09/2025 | Negative | B1 | 31 | 210 | 0 | 1 | 0 | -1 |
| Bird bath / rain water | 26/09/2025 | Negative | B2 | 30.1 | 208 | 0 | 0 | 0 | -1 |
| Bird bath / rain water | 20/09/2025 | Negative | B1 | 32.2 | 178 | 0 | 1 | 0 | -1 |
| Bird bath / rain water | 17/09/2025 | Negative | B2 | 31.3 | 195 | 0 | -1 | 0 | -1 |
| Bird bath / rain water | 16/09/2025 | Negative | B2 | 30.5 | 225 | 0 | -1 | 0 | 0 |
| Bird bath / rain water | 13/09/2025 | Negative | B2 | 31.3 | 197 | 0 | -1 | 0 | 0 |
| Bird bath / rain water | 11/09/2025 | Negative | B1 | 30.4 | 214 | 0 | 4 | 0 | 1 |
| Bird bath / rain water | 04/09/2025 | Negative | B2 | 30.7 | 209 | 0 | 0 | 0 | -1 |
| Bird bath / rain water | 02/09/2025 | Negative | B2 | 30.7 | 214 | 0 | -1 | 0 | -1 |
| Bird bath / rain water | 31/08/2025 | Negative | B2 | 31.8 | 174 | 0 | -1 | 0 | 0 |
| Bird bath / rain water | 29/08/2025 | Negative | B2 | 30.1 | 231 | 0 | 0 | 0 | -1 |
| Bird bath / rain water | 28/08/2025 | Negative | B2 | 31.4 | 190 | 0 | -1 | 0 | -1 |
| Bird bath / rain water | 26/08/2025 | Negative | B2 | 31.1 | 193 | 0 | -1 | 0 | 0 |
| Bird bath / rain water | 25/08/2025 | Negative | B2 | 31.3 | 183 | 0 | -1 | 0 | 0 |
| Bird bath / rain water | 22/08/2025 | Negative | B2 | 30.9 | 220 | 0 | 0 | 0 | 0 |
| Bird bath / rain water | 21/08/2025 | Negative | B1 | 30.7 | 214 | 0 | 1 | 0 | -1 |
| Bird bath / rain water | 21/08/2025 | Negative | B2 | 30.3 | 204 | 0 | -3 | 0 | 1 |
| Bird bath / rain water | 18/08/2025 | Negative | B1 | 31 | 208 | 0 | 1 | 0 | -1 |
| Bird bath / rain water | 25/07/2025 | Negative | B2 | 31.3 | 194 | 0 | -1 | 0 | 0 |
| Bird bath / rain water |  | Negative | B2 | 33.1 | 140 | 0 | 0 | 0 | 1 |
| Bird bath / rain water |  | Negative | B1 | 31.1 | 184 | 0 | -1 | 0 | -1 |
