## Supplementary Table 5 for "Rapid GeneXpert surveillance of influenza A virus in seabirds and the environment provides early warning for wildlife health in Aotearoa New Zealand"

**Supplementary Table 5**. Ct and end-point values with environmental samples collected from an urban duck pond in the Dunedin Botanic Gardens. Sample types inclided frsh duck faeces from mallard ducks, passive filtered water, water and sediment.

| **Ct** | | | | | |  | **End point** | | | | | |
| --- | --- | --- | --- | --- | --- | --- | --- | --- | --- | --- | --- | --- |
| Sample type | FluA1 | FluA2 | FluB | RSV | SPC |  | Sample type | FluA1 | FluA2 | FluB | RSV | SPC |
| Faeces | 0 | 0 | 0 | 0 | 29.8 |  | Faeces | 0 | 0 | 0 | 0 | 232.0 |
| Faeces | 0 | 0 | 0 | 0 | 30.6 |  | Faeces | 0 | 0 | 0 | 0 | 139.0 |
| Faeces | 0 | 0 | 0 | 0 | 32.1 |  | Faeces | 0 | 0 | 0 | 0 | 227.0 |
| Faeces | 0 | 0 | 0 | 0 | 32.8 |  | Faeces | 0 | 0 | 0 | 0 | 142.0 |
| Faeces | 0 | 0 | 0 | 0 | 30.7 |  | Faeces | 0 | 0 | 0 | 0 | 168.0 |
| Faeces | 0 | 0 | 0 | 0 | 33.6 |  | Faeces | 0 | 0 | 0 | 0 | 112.0 |
| Passive filter | 0.0 | 0.0 | 0.0 | 0.0 | 33.0 |  | Passive filter | 1.0 | 0.0 | -1.0 | -1.0 | 130.0 |
| Passive filter | 0.0 | 0.0 | 0.0 | 0.0 | 31.3 |  | Passive filter | 1.0 | 0.0 | 0.0 | 0.0 | 176.0 |
| Passive filter | 0.0 | 0.0 | 0.0 | 0.0 | 34.0 |  | Passive filter | 1.0 | 12.0 | 0.0 | -1.0 | 131.0 |
| Passive filter | 0.0 | 0.0 | 0.0 | 0.0 | 30.4 |  | Passive filter | 1.0 | -1.0 | 0.0 | -1.0 | 226.0 |
| Passive filter | 0.0 | 0.0 | 0.0 | 0.0 | 30.2 |  | Passive filter | 2.0 | 0.0 | 0.0 | -2.0 | 232.0 |
| Passive filter | 0.0 | 0.0 | 0.0 | 0.0 | 31.6 |  | Passive filter | 5.0 | -1.0 | -1.0 | -1.0 | 179.0 |
| Passive filter | 0.0 | 0.0 | 0.0 | 0.0 | 32.5 |  | Passive filter | 1.0 | 5.0 | -1.0 | 3.0 | 135.0 |
| Passive filter | 0.0 | 0.0 | 0.0 | 0.0 | 30.5 |  | Passive filter | 2.0 | 0.0 | -1.0 | 0.0 | 199.0 |
| Passive filter | 0.0 | 0.0 | 0.0 | 0.0 | 31.7 |  | Passive filter | -1.0 | -1.0 | 0.0 | -2.0 | 169.0 |
| Passive filter | 0.0 | 0.0 | 0.0 | 0.0 | 30.1 |  | Passive filter | 2.0 | 0.0 | 0.0 | -1.0 | 226.0 |
| Passive filter | 0.0 | 0.0 | 0.0 | 0.0 | 29.6 |  | Passive filter | 15.0 | 0.0 | -1.0 | -1.0 | 227.0 |
| Passive filter | 0.0 | 0.0 | 0.0 | 0.0 | 30.9 |  | Passive filter | 2.0 | -1.0 | 0.0 | -1.0 | 215.0 |
| Passive filter | 0.0 | 0.0 | 0.0 | 0.0 | 29.8 |  | Passive filter | 2.0 | 0.0 | 0.0 | 0.0 | 209.0 |
| Passive filter | 0.0 | 0.0 | 0.0 | 0.0 | 29.8 |  | Passive filter | 2.0 | 0.0 | 0.0 | 0.0 | 209.0 |
| Water | 0.0 | 0.0 | 0.0 | 0.0 | 31.4 |  | Water | 1.0 | 2.0 | -1.0 | 0.0 | 180.0 |
| Water | 0.0 | 0.0 | 0.0 | 0.0 | 31.5 |  | Water | 12.0 | 0.0 | 0.0 | 0.0 | 185.0 |
| Water | 0.0 | 0.0 | 0.0 | 0.0 | 32.5 |  | Water | 28.0 | 1.0 | -1.0 | 0.0 | 161.0 |
| Water | 0.0 | 0.0 | 0.0 | 0.0 | 30.2 |  | Water | 29.0 | 0.0 | 0.0 | 0.0 | 196.0 |
| Water | 0.0 | 0.0 | 0.0 | 0.0 | 30.5 |  | Water | 12.0 | 6.0 | 0.0 | 0.0 | 200.0 |
| Water | 34.2 | 0.0 | 0.0 | 0.0 | 30.9 |  | Water | 210.0 | 45.0 | -1.0 | 0.0 | 151.0 |
| Water | 35.2 | 0.0 | 0.0 | 0.0 | 31.1 |  | Water | 117.0 | 40.0 | -3.0 | -1.0 | 140.0 |
| Water | 35.5 | 0.0 | 0.0 | 0.0 | 30.7 |  | Water | 106.0 | 16.0 | -1.0 | 1.0 | 143.0 |
| Water | 34.7 | 0.0 | 0.0 | 0.0 | 30.2 |  | Water | 133.0 | 23.0 | -3.0 | 0.0 | 181.0 |
| Water | 34.4 | 37.9 | 0.0 | 0.0 | 30.2 |  | Water | 171.0 | 53.0 | 0.0 | 1.0 | 147.0 |
| Water | 37.2 | 0.0 | 0.0 | 0.0 | 32.7 |  | Water | 101.0 | 25.0 | -1.0 | 1.0 | 152.0 |
| Water | 34.2 | 36.7 | 0.0 | 0.0 | 32.5 |  | Water | 311.0 | 91.0 | -3.0 | 1.0 | 161.0 |
| Water | 38.7 | 0.0 | 0.0 | 0.0 | 37.6 |  | Water | 77.0 | 0.0 | -4.0 | 2.0 | 57.0 |
| Water | 34.5 | 37.8 | 0.0 | 0.0 | 32.0 |  | Water | 338.0 | 92.0 | -6.0 | 0.0 | 153.0 |
| Water | 35.8 | 39.3 | 0.0 | 0.0 | 33.7 |  | Water | 201.0 | 61.0 | -4.0 | 1.0 | 129.0 |
| Sediment | 0.0 | 0.0 | 0.0 | 0.0 | 30.6 |  | Sediment | -1.0 | 0.0 | -1.0 | 0.0 | 176.0 |
| Sediment | 0.0 | 0.0 | 0.0 | 0.0 | 31.1 |  | Sediment | -3.0 | -1.0 | -1.0 | 1.0 | 187.0 |
| Sediment | 0.0 | 0.0 | 0.0 | 0.0 | 29.8 |  | Sediment | 2.0 | 0.0 | 0.0 | 0.0 | 220.0 |
| Sediment | 0.0 | 0.0 | 0.0 | 0.0 | 31.1 |  | Sediment | 1.0 | 0.0 | 0.0 | -1.0 | 204.0 |
| Sediment | 0.0 | 0.0 | 0.0 | 0.0 | 30.7 |  | Sediment | 1.0 | 0.0 | 0.0 | -1.0 | 173.0 |
| Sediment | 0.0 | 0.0 | 0.0 | 0.0 | 30.4 |  | Sediment | 1.0 | 0.0 | 0.0 | 0.0 | 216.0 |
| Sediment | 0.0 | 0.0 | 0.0 | 0.0 | 30.5 |  | Sediment | 23.0 | 1.0 | -2.0 | -1.0 | 198.0 |
| Sediment | 0.0 | 0.0 | 0.0 | 0.0 | 30.7 |  | Sediment | 2.0 | 0.0 | 1.0 | 0.0 | 190.0 |
| Sediment | 0.0 | 0.0 | 0.0 | 0.0 | 30.7 |  | Sediment | 20.0 | -1.0 | -1.0 | 0.0 | 203.0 |
| Sediment | 0.0 | 0.0 | 0.0 | 0.0 | 30.2 |  | Sediment | 0.0 | -1.0 | 0.0 | -1.0 | 185.0 |
| Sediment | 0.0 | 0.0 | 0.0 | 0.0 | 32.4 |  | Sediment | 11.0 | 0.0 | -1.0 | -1.0 | 179.0 |
| Sediment | 0.0 | 0.0 | 0.0 | 0.0 | 34.0 |  | Sediment | 1.0 | 0.0 | 0.0 | 0.0 | 159.0 |
| Sediment | 0.0 | 0.0 | 0.0 | 0.0 | 31.2 |  | Sediment | 15.0 | -1.0 | 0.0 | 1.0 | 193.0 |
| Sediment | 0.0 | 0.0 | 0.0 | 0.0 | 31.5 |  | Sediment | 9.0 | 12.0 | 0.0 | -1.0 | 165.0 |
| Sediment | 0.0 | 0.0 | 0.0 | 0.0 | 31.3 |  | Sediment | 1.0 | 0.0 | 0.0 | 0.0 | 176.0 |
