## Supplementary Table 6 for "Rapid GeneXpert surveillance of influenza A virus in seabirds and the environment provides early warning for wildlife health in Aotearoa New Zealand"

**Supplementary Table 6**. Error log for the GeneXpert while deployed. No ERRORS observed prior to transfer to Albatross Centre.

| Sample ID | Notes | Run Call | Status | Mod | Error Code and descriptor |
| --- | --- | --- | --- | --- | --- |
| DBG.FA.RNA005 | DBG.Feeding Area.RNA Shield | ERROR | Aborted | B2 | Operation terminated Error 2125: Termination Error - Insufficient Volume: 14, 30, 0, 0 |
| DBG.FA.RNA004 | DBG.Feeding Area.RNA Shield | INVALID | Done | B1 | Probe Checks all passed. SPC failed |
| DBG.FA.RNA003 | DBG.Feeding Area.RNA Shield | ERROR | Aborted | B2 | Operation terminated Error 2125: Termination Error - Insufficient Volume: 14, 30, 0, 0 |
| DBG.FA.PBS004 | DBG.Feeding Area.PBS | ERROR | Aborted | B2 | Operation terminated, Error 2005: Motion of the syringe drive was not detected. Detected motion started at position 32 uL and transferred 395 uL at valve position 491 with pressure -4.8 PSI |
| DBG.FA.PF002 | DBG.Feeding Area.Passive Filter | No Result | Stopped | B2 | Operator |
| DBG.CL.PF005 | DBG.Clive Lister.Passive Filter | No Result | Aborted | B1 | Operation terminated. Error 2037: The cartridge integrity test failed at valve position 1390. The pressure change of 0.6 PSI did not exceed the requirement of 4.0 PSI. The pressure increased from 3.7 PSI to 4.4 PSI during the test |
| Error test 1 | Tap Water Rat room | ERROR | Incomplete | B2 | Probe Checks all passed. [Communication Lost Code 2123: Module B2 lost communication while test was running, attempting recovery]; [Operation terminated. Error 2126: Module B2 was reset]; [Communication Restored. Code 2124: Module B2 communication restored] |
| Test2 | Tap Water Rat room | ERROR | Incomplete | B2 | Probe Checks all passed. Operation terminated. Error 2126: Module B2 was reset |
| water15oct2025 | water15oct2025 | ERROR | Aborted | B1 | Operation terminated. Error 2008: Syringe pressure reading of 100.1 PSI exceeds the protocol limit of 100.0 PSI |
| water bath sample 2-10-25 | water bath sample 2-10-25 | ERROR | Aborted | B1 | Probe Check Flu B failed. Post-run analysis error. Error 5007: [Flu B] probe check failed. Probe check value of 103 for reading number 2 was below the minimum of 104 |
| Guano top carpark | Guano top carpark | ERROR | Aborted | B1 | Operation terminated. Error 2008: Syringe pressure reading of 100.1 PSI exceeds the protocol limit of 100.0 PSI |
| guanosept13th | guanosept13th | ERROR | Aborted | B1 | Operation terminated. Error 2008: Syringe pressure reading of 100.1 PSI exceeds the protocol limit of 100.0 PSI |
| bird bath RAC staff park 12thSep | bird bath RAC staff park 12thSep | ERROR | Aborted | B1 | Operation terminated. Error 2008: Syringe pressure reading of 100.1 PSI exceeds the protocol limit of 100.0 PSI |
| Bath 09-09-25 | Bath 09-09-25 | ERROR | Aborted | B1 | Operation terminated. Error 2008: Syringe pressure reading of 100.1 PSI exceeds the protocol limit of 100.0 PSI |
| bath 08-09-2025 | bath 08-09-2025 | ERROR | Aborted | B1 | Operation terminated. Error 2008: Syringe pressure reading of 100.1 PSI exceeds the protocol limit of 100.0 PSI |
| birdbath#2 | birdbath#2 | ERROR | Aborted | B1 | Operation terminated. Error 2008: Syringe pressure reading of 100.1 PSI exceeds the protocol limit of 100.0 PSI |
| Bird poo sample 04th Sep | Bird poo sample 04th Sep | ERROR | Aborted | B1 | Operation terminated. Error 2008: Syringe pressure reading of 100.1 PSI exceeds the protocol limit of 100.0 PSI |
| bathwater B1 26-08-25 | bathwater B1 26-08-25 | ERROR | Aborted | B1 | Operation terminated. Error 2008: Syringe pressure reading of 100.1 PSI exceeds the protocol limit of 100.0 PSI |
| Bird Bath OBS#1 25 Aug | Bird Bath OBS#1 25 Aug | ERROR | Aborted | B1 | Operation terminated. Error 2008: Syringe pressure reading of 100.1 PSI exceeds the protocol limit of 100.0 PSI |
| rain water sample B. 22-Aug-25 | rain water sample B. 22-Aug-25 | ERROR | Aborted | B1 | Operation terminated. Error 2008: Syringe pressure reading of 100.1 PSI exceeds the protocol limit of 100.0 PSI |
| water sample 18.Aug tap water | water sample 18.Aug tap water | ERROR | Aborted | B2 | Operation terminated. Error 2008: Syringe pressure reading of 100.1 PSI exceeds the protocol limit of 100.0 PSI |
| guano test 2. 13-8-25 | guano test 2. 13-8-25 | ERROR | Aborted | B1 | Operation terminated. Error 2008: Syringe pressure reading of 100.1 PSI exceeds the protocol limit of 100.0 PSI |
| guano test 13825 | guano test 13825 | ERROR | Aborted | B2 | Operation terminated. Error 2008: Syringe pressure reading of 100.1 PSI exceeds the protocol limit of 100.0 PSI |
| guano 07-08-2025 (B1) | guano 07-08-2025 (B1) | ERROR | Aborted | B1 | Operation terminated. Error 2008: Syringe pressure reading of 100.1 PSI exceeds the protocol limit of 100.0 PSI |
| water 25-07-25 | water 25-07-25 | ERROR | Aborted | B1 | Operation terminated. Error 2008: Syringe pressure reading of 100.1 PSI exceeds the protocol limit of 100.0 PSI |
| guano 17-07-2025 | guano 17-07-2025 | ERROR | Aborted | B1 | Operation terminated. Error 2008: Syringe pressure reading of 100.1 PSI exceeds the protocol limit of 100.0 PSI |
| water71525 | water71525 | ERROR | Aborted | B2 | Operation terminated. Error 2008: Syringe pressure reading of 100.1 PSI exceeds the protocol limit of 100.0 PSI |
| bath 6-23-25 (1) | bath 6-23-25 | ERROR | Aborted | B2 | Operation terminated. Error 2008: Syringe pressure reading of 100.1 PSI exceeds the protocol limit of 100.0 PSI |
| guano 6-23-25 | guano 6-23-25 | ERROR | Aborted | B1 | guano from car park: Operation terminated. Error 2008: Syringe pressure reading of 100.1 PSI exceeds the protocol limit of 100.0 PSI |
| guano 2 | guano 2 | ERROR | Aborted | B1 | Operation terminated. Error 2008: Syringe pressure reading of 100.1 PSI exceeds the protocol limit of 100.0 PSI: guano sample outside of obs |
